# *Lactobacillus* Restructures the Micro/Mycobiome to Combat Glycoprotein-130 Associated Microtubule Remodeling and Right Ventricular Dysfunction in Pulmonary Arterial Hypertension

**DOI:** 10.1101/2024.08.19.608469

**Authors:** Sasha Z. Prisco, Madelyn Blake, Felipe Kazmirczak, Ryan Moon, Neal Vogel, Daphne Moutsoglou, Thenappan Thenappan, Kurt W. Prins

## Abstract

Emerging data demonstrate systemic and local inflammation regulate right ventricular (RV) adaption in preclinical and human pulmonary arterial hypertension (PAH). Pathological RV inflammation is targetable as antagonism of glycoprotein-130 (GP130) signaling counteracts pathological microtubule remodeling and improves RV function in rodents. Microtubules control several aspects of cardiomyocyte biology including cellular and nuclear size/structure, t-tubule homeostasis, and the proper localization of connexin-43. The intestinal microbiome regulates systemic inflammation, but the impact of the gut microbiome on the GP130-microtubule axis in RV failure is unknown. Here, we examined how the anti-inflammatory bacteria, *Lactobacillus*, modulated cellular and physiological RV phenotypes in preclinical and clinical PAH. *Lactobacillus* supplementation restructured the gut micro/mycobiome, suppressed systemic inflammation, combatted pathological GP130-mediated RV cardiomyocyte microtubule remodeling, and augmented RV function in rodent PAH. Moreover, *Lactobacillus* was associated with superior RV adaption in human PAH. These data further support the hypothesis that inflammation negatively impacts RV adaption in PAH, and identify the gut microbiome as a potentially targetable regulator of RV function in PAH.

## Introduction

Emerging data demonstrate systemic and local inflammation regulate right ventricular (RV) adaption in preclinical and human pulmonary arterial hypertension (PAH)^1^. Pathological RV inflammation is targetable as antagonism of glycoprotein-130 (GP130) signaling counteracts pathological microtubule remodeling and improves RV function in rodents^2^. Microtubules control several aspects of cardiomyocyte biology including cellular and nuclear size/structure, t-tubule homeostasis, and the proper localization of connexin-43^3^. Thus, restoration of microtubule regulation following cardiac stress is cardioprotective^3^. The intestinal microbiome regulates systemic inflammation^4^, but the impact of the gut microbiome on the GP130-microtubule axis in RV failure is unknown. Here, we examined how the anti-inflammatory bacteria, *Lactobacillus*^*4*^, modulated cellular and physiological RV phenotypes in preclinical and clinical PAH.

## Methods

Male Sprague Dawley rats (200-250 g) received a subcutaneous injection of 60 mg/kg monocrotaline (MCT) or phosphate-buffered saline (control). Two weeks following MCT injection, rats were randomly allocated to daily *Lactobacillus rhamnosus* (4×10^7^ colony-forming units, Chr. Hansen) in one mL sterile water (MCT-*Lactobacillus*) or one mL sterile water (MCT-Water) via oral gavage for ten days. Next-generation metagenomics and internal transcribed spacer 2 sequencing evaluated the fecal composition of the bacteria and fungi, respectively. SomaScan proteomics quantified the serum proteome. KEGG pathway analysis (ShinyGO 0.80) delineated pathways altered in the serum proteome. Immunoblots of RV protein extracts (25 μg) using antibodies to α-tubulin (Sigma-Aldrich:T6199), β-tubulin (Sigma-Aldrich:T4026), and detyrosinated α-tubulin (Abcam:ab48389) were performed using the Odyssey Infrared Imaging system^2^. RV and jejunal formalin sections were subjected to heat-mediated antigen retrieval (Reveal Decloaker, Biocare Medical), stained with primary antibodies to β-tubulin (Abcam:ab6046), connexin-43 (Abcam:ab235585), and then AlexaFluor-568 anti-rabbit secondary antibody (ThermoScientific:A-11036), Wheat Germ Agglutinin

AlexaFluor-633 (ThermoFisher:W21404), and AlexaFluor-488 Phalloidin (AAT Bioquest:23153). Confocal micrographs were collected on a Zeiss LSM900 Airyscan 2.0 confocal microscope. Confocal and histological images were blindly analyzed by MB, RM, and NV using FIJI. Echocardiography and invasive closed-chest pressure-volume loops^2^ evaluated rodent PAH severity and RV function. All animal studies were approved by the UMN Institutional Animal Care and Use Committee. Statistical analyses were completed with GraphPad Prism 10.1 and MetaboAnalyst software (https://www.metaboanalyst.ca/).

## Results

*Lactobacillus* supplementation restructured the intestinal bacterial ecosystem as MCT-*Lactobacillus* rats exhibited a microbiome composition that was an intermediate between control and MCT-Water (**Figure A**). Variable of importance in projection scores identified *Lactobacillus* as the most important bacteria for differentiating the three groups. *Lactobacillus* gavage increased the abundance of *Lactobacillus* as compared to MCT-Water, but it was not fully restored. In addition, *Lactobacillus* supplementation altered the mycobiome (**Figure A**). The mycobiome change was not as distinct as the microbiome in the experimental groups, however MCT-*Lactobacillus* more closely mirrored control than MCT-Water. Finally, correlational analysis identified multiple bacteria and fungi that were either positively or negatively associated with *Lactobacillus* abundance (**Figure A**).

**Figure:**
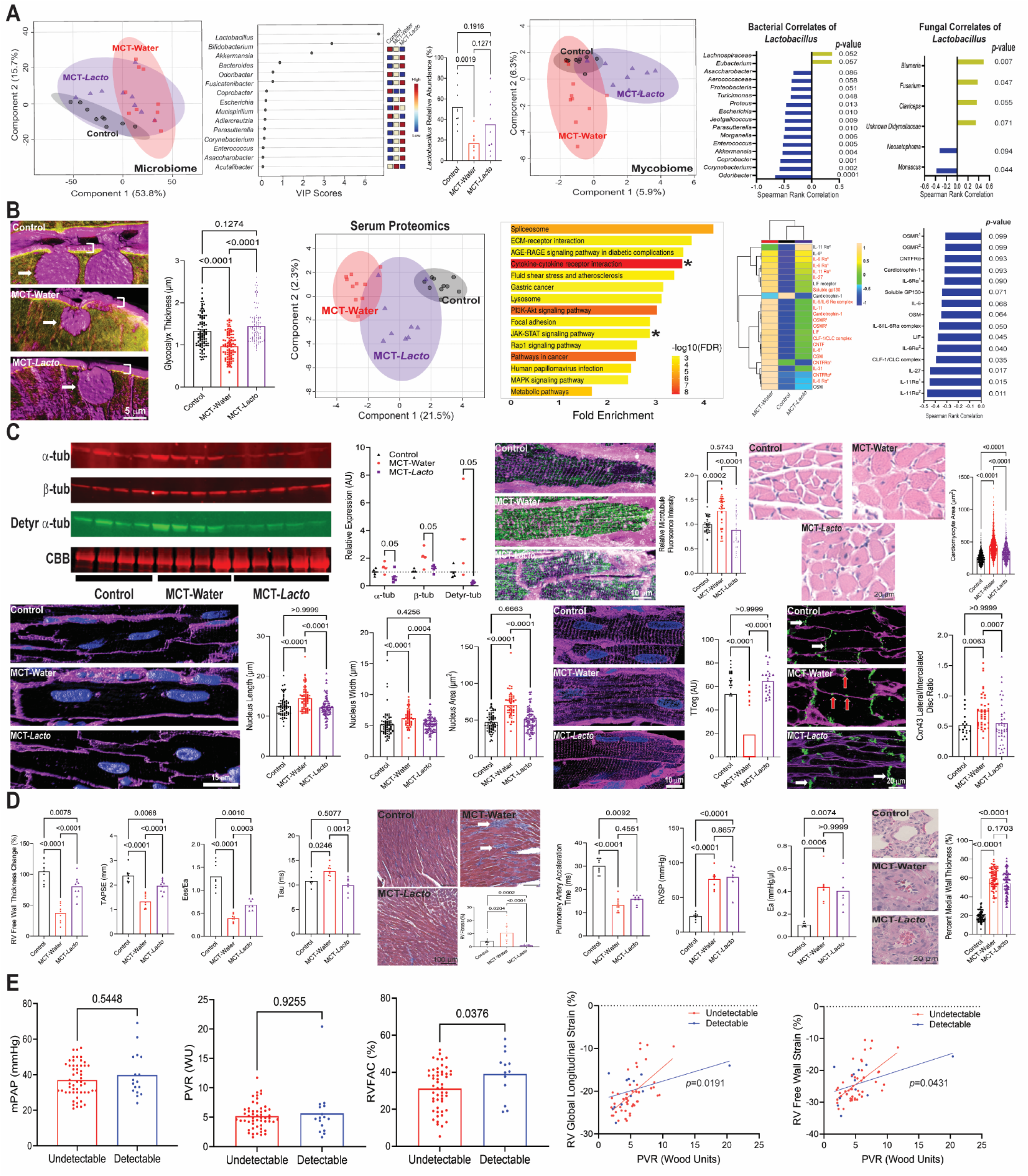
*Lactobacillus* supplementation restructures the intestinal microbiome/mycobiome, suppresses pathological inflammation, mitigates pathological GP130-mediated microtubule remodeling, and enhances right ventricular function in pulmonary arterial hypertension. (**A**) Partial least squares discriminant analysis revealed *Lactobacillus* supplementation restructured the gut microbiome to a composition intermediate of control and MCT-Water. Variable of importance in projection scores defined the fifteen bacterial species that most distinguished the intestinal microbiome of control, MCT-Water, and MCT-*Lactobacillus*, and *Lactobacillus* was the most important. *Lactobacillus* gavage increased the abundance of *Lactobacillus* as compared to MCT-Water, but did not restore to control levels (*p*-values determined by one-way ANOVA with Tukey’s multiple comparisons test). Evaluation of the fecal mycobiome by partial least squares discriminant analysis. Spearman rank correlation determined the bacteria and fungi either positively or negatively correlated with *Lactobacillus*. (**B**) Representative super resolution confocal micrographs of jejunum evaluated glycocalyx thickness (bracket). *Lactobacillus* supplemented restored glycocalyx thickness (*p*-values determined by Kruskal-Wallis test with Dunn’s multiple comparison test). Arrows delineate goblet cells. Sparse partial least squares discriminant analysis revealed *Lactobacillus* altered the serum proteome. KEGG pathway analysis of differentially regulated proteins identified the cytokine-cytokine receptor interaction and JAK-STAT signaling pathways (denoted by asterisks). Profiling of GP130 ligands with hierarchical cluster analysis showed *Lactobacillus* decreased all serum GP130 ligands compared to MCT-Water except for one of the IL-11 receptor α aptamers (ligands in red text are statistically different between groups as determined by one-way ANOVA with Tukey’s multiple comparison test if normally distributed or Kruskal-Wallis test with Dunn’s multiple comparisons test if not normally distributed). Spearman rank correlation determined the GP130 ligands that correlated with *Lactobacillus* abundance. (**C**) Representative immunoblots of RV tissue demonstrated *Lactobacillus* supplementation mitigated pathological microtubule remodeling by decreasing α-tubulin, β-tubulin, and detyrosinated α-tubulin abundance as compared to MCT-Water (*p*-values determined by unpaired *t-*test using control as an arbitrary value of 1). Western blot results were normalized to the myosin heavy chain band in the Coomassie brilliant blue (CBB) stained post-transfer gel. Representative super resolution confocal micrographs of RV sections revealed *Lactobacillus* decreased microtubule density (green: β-tubulin, purple: wheat germ agglutinin [WGA]) (*p*-value assessed by Kruskal-Wallis test with Dunn’s multiple comparison test). Representative histological images of RV tissue showed *Lactobacillus* dampened RV cardiomyocyte hypertrophy (*p*-value determined by Kruskal-Wallis test with Dunn’s multiple comparison test). Representative confocal microcytes of RV cardiomyocytes demonstrated *Lactobacillus* normalized nuclear length, width, and area (blue: DAPI, purple: WGA) (*p*-values determined by Kruskal-Wallis test with Dunn’s multiple comparison test). Representative confocal micrographs of RV cardiomyocytes showed *Lactobacillus* supplementation restored the normal striated t-tubule architecture (*p*-values determined by Kruskal-Wallis test with Dunn’s multiple comparison test). *Lactobacillus* supplementation prevented lateralization of connexin-43 (white arrows: connexin-43 at gap junction, red arrows: lateralized connexin-43) (*p*-values determined by Kruskal-Wallis test with Dunn’s multiple comparison test). (**D**) *Lactobacillus* supplementation augmented RV systolic function as assessed by RV free wall thickness change, tricuspid annular plane systolic excursion (TAPSE), and RV-pulmonary artery coupling determined by the ratio of end-systolic elastance to effective arterial elastance (Ees/Ea) and improved RV diastolic function (tau). *Lactobacillus* supplementation decreased RV fibrosis (arrows delineate areas of fibrosis). *Lactobacillus* supplementation did not significantly change pulmonary artery acceleration time, right ventricular systolic pressure (RVSP), Ea, or pulmonary arteriole remodeling as assessed histologically. *P*-values determined by one-way ANOVA with Tukey’s multiple comparison test for RV free wall thickness change, TAPSE, RVSP, and tau; Kruskal-Wallis test with Dunn’s multiple comparison test for pulmonary artery acceleration time, Ea, and percent medial wall thickness; and Brown-Forsythe ANOVA test with Dunnett’s T3 multiple comparisons test for Ees/Ea as there was unequal variance. (**E**) *Lactobacillus* abundance was not associated with differences in mean pulmonary arterial pressure (mPAP) and pulmonary vascular resistance (PVR) in PAH patients. PAH patients with detectable *Lactobacillus* had numerically higher RV fractional area change (RVFAC) and superior RV adaptability to increased afterload as demonstrated by the relationships between RV global longitudinal strain or RV free wall strain to PVR. *P*-values determined by Mann-Whitney test for mPAP and PVR, unpaired t-test for RVFAC, and simple linear regression (statistical difference in slope) when determining the relationship between RV global/free wall strain and PVR.

Next, we evaluated how *Lactobacillus*-mediated restructuring of the micro/mycobiome impacted the intestinal barrier and systemic inflammation. Confocal microscopy demonstrated *Lactobacillus* restored jejunal glycocalyx thickness, which would improve intestinal barrier function (**Figure B**). To evaluate this hypothesis, we used SomaScan-based proteomics to probe the systemic inflammatory response. Sparse partial least squares discriminant analysis revealed *Lactobacillus* shifted the serum proteome towards control (**Figure B**). KEGG pathway analysis of differentially regulated serum proteins identified cytokine-cytokine receptor interaction and JAK-STAT signaling pathways. To explore GP130 signaling specifically, we profiled serum GP130 ligands/signaling molecules and found *Lactobacillus* decreased almost all of the serum GP130 signaling molecules compared to MCT-Water, suggesting *Lactobacillus* dampened systemic inflammation. Importantly, *Lactobacillus* abundance was inversely correlated with all GP130 cytokines/signaling molecules (**Figure B**), lending further support that *Lactobacillus* imparted an anti-inflammatory effect via GP130 signaling.

Then, we evaluated how dampened GP130 signaling altered the RV cardiomyocyte microtubule phenotype. Immunoblots and confocal microscopy demonstrated *Lactobacillus* suppressed microtubule stabilization in RV cardiomyocytes (**C**). Moreover, *Lactobacillus* counteracted cardiomyocyte and nuclear hypertrophy, restored t-tubule architecture, and prevented mislocalization of connexin-43 (**C**).

To probe the physiological consequences of the observed cellular microtubule phenotypes, echocardiography and invasive pressure-volume loop analysis quantified RV function. *Lactobacillus* supplementation improved echocardiographic and hemodynamic measures of systolic and diastolic parameters of RV function and reduced RV fibrosis independent of afterload (**D**).

To provide human relevance to our preclinical findings, we determined how fecal *Lactobacillus* abundance determined by metagenomics sequencing was associated with RV adaptability in our previously evaluated PAH population^5^. Mean pulmonary arterial pressure/pulmonary vascular resistance were not different when patients were dichotomized based on the presence/absence of *Lactobacillus* (**E**). However, PAH patients with detectable *Lactobacillus* had higher RV fractional area change and superior RV adaptability to increasing afterload (**E**).

## Discussion

In summation, *Lactobacillus* supplementation restructured the gut micro/mycobiome, suppressed systemic inflammation, combatted pathological GP130-mediated RV cardiomyocyte microtubule remodeling, and augmented RV function in rodent PAH. Moreover, *Lactobacillus* was associated with superior RV adaption in human PAH. These data further support the hypothesis that inflammation negatively impacts RV adaption in PAH, and identify the gut microbiome as a potentially targetable regulator of RV function in PAH.

